# *Trypanosoma cruzi* amastigotes have a reduced replication rate during chronic stage infections

**DOI:** 10.1101/2020.08.04.236240

**Authors:** Alexander I. Ward, Francisco Olmo, Richard Atherton, Martin C. Taylor, John M. Kelly

**Affiliations:** Department of Infection Biology, London School of Hygiene and Tropical Medicine, London, UK

## Abstract

Chronic *Trypanosoma cruzi* infections are typically life-long, with small numbers of parasites surviving in restricted tissue sites, which include the gastro-intestinal tract. There is considerable debate about the replicative status of these persistent parasites. Here, we investigated *T. cruzi* proliferation in the colon of chronically infected mice using 5-ethynyl-2’deoxyuridine incorporation into DNA to provide “snapshots” of parasite replication. Highly sensitive imaging of infection foci at single cell resolution revealed that parasites are three times more likely to be in S-phase during the acute stage than during the chronic stage. By implication, chronic infections are associated with a reduced rate of parasite replication. Despite this, very few host cells survive infection for >14 days, suggesting that *T. cruzi* persistence continues to involve regular cycles of replication, host cell lysis and re-infection. Therefore, long-term persistence in the colon is more likely to be associated with reduced proliferation than with dormancy.

## Introduction

Disease latency, mediated by a wide range of mechanisms, is a common feature of viral, bacterial and parasitic infections (Weidner-Glunde *et al*. 2020; Dodd and Schlesinger, 2017; Barrett *et al*. 2019). However, there can be long-term consequences for the host, which include relapse and/or inflammatory pathology. The terms “persistent”, “dormant” and “metabolically quiescent” are used, often interchangeably, to describe pathogens in this state. The phenomenon has evolved independently and frequently in different pathogen groups, presumably because it acts to enhance survival and transmission. The “persister” phenotype does not involve the acquisition of selected mutations, and is often associated with treatment failure, antibiotic tolerance being the best studied example (Pontes *et al*. 2020; Mandal *et al*. 2019). In the case of Chagas disease, some form of dormancy or restricted replication has been widely postulated as a mechanism that might explain long-term parasite survival and the high rate of treatment failure (Francisco *et al*. 2017).

Chagas disease is caused by the protozoan parasite *Trypanosoma cruzi*, which infects 6-7 million people, mainly in Latin America. Better drugs and innovative immunological interventions are urgently required. Human infection is normally initiated when faeces of the triatomine insect vector, contaminated with metacyclic trypomastigote forms of the parasite, come into contact with the bite wound, or when they are rubbed into mucous membranes. An acute parasitaemia develops, which can be asymptomatic, or manifest as generalised symptoms such as fever, headache and muscle pain. Suppression of the infection is then mediated by a CD8+ T cell mediated response which reduces parasite numbers to extremely low levels (Cardillo *et al.* 2015, Tarleton, 2015). A subset of infected individuals (~30%) eventually develop the classical Chagasic cardiac and digestive symptoms, although this can be decades after the acute stage infection. Dilated cardiomyopathy and digestive megasyndromes are the most common morbidities, and can often be fatal (Bonney *et al*. 2019; Martinez *et al*. 2020). It remains to be established how the parasite is able to persist long-term, albeit at very low levels, in the face of a robust adaptive immune response (Pack *et al*. 2018). Furthermore, the reasons why treatment failures are a common outcome need to be better understood at a mechanistic level to guide the design of improved chemotherapy (Gaspar *et al*. 2015). In this context, the recent report of a non-proliferative form of *T. cruzi* that is refractory to treatment with the front-line drug benznidazole (Sánchez-Valdéz *et al*. 2018) could have important implications.

The ability of human parasites to enter a long-term quiescent state, in which both replication and metabolism are slowed, has been described in *Toxoplasma gondii* (the bradyzoite) (Krishnan *et al.* 2020), and some *Plasmodium* species (Barrett *et al*. 2019). As with many prokaryotic pathogens (Fisher *et al*. 2017; Gollan *et al*. 2019), the ‘dormant’ state involves lower levels of DNA synthesis and transcription, down-regulation of energy catabolism, and activation of DNA damage/cellular stress responses. In *T. gondii*, a master transcription factor (BFD1), activated by stress response pathways, initiates the on-set of bradyzoite development (Waldman *et al*. 2020). The precise triggers that lead to differentiation into the quiescent hypnozoite liver stage in some *Plasmodium spp.* have been elusive (Barrett *et al*. 2019). Amongst eukaryotic pathogens, these examples represent one end of the “dormancy spectrum”, in which entry into a quiescent metabolic state for extended periods. It has also been tentatively proposed that *Leishmania donovani* can undergo a form of dormancy, although the mechanisms involved are unknown (Tegazzini *et al*. 2016). The situation in other *Leishmania spp.* is more definitive, with the identification of partially quiescent intracellular amastigotes which exhibit a slower metabolic flux and a reduced replication rate (Kloehn *et al*. 2015; Mandell and Beverley, 2017). This stops short of full long-term dormancy in which parasites enter G0/G1 cell cycle arrest. *Plasmodium falciparum* blood stage schizonts can also enter a transient state of dormancy, induced by treatment with the front-line drug artemisinin (Tucker *et al*. 2012; Teuscher *et al*. 2012). This capacity to respond to stress by halting progress through the cell cycle exists in most cells that have DNA damage sensing machinery (Lanz *et al*. 2019; Verma *et al*. 2019). The existence of a non-replicative phenotype in mammalian forms of the African trypanosome, *Trypanosoma brucei*, beyond the G0/G1 arrested stumpy form required for onward transmission (Silvester *et al*. 2017), remains speculative.

Observations of both *in vitro* and *in vivo T. cruzi* infections identified a sub-population of non-dividing intracellular amastigotes that retained the ability to differentiate into flagellated trypomastigotes, which were then able to propagate the infection (Sánchez-Valdéz *et al*. 2018). This phenomenon was defined as spontaneous dormancy on the basis of experiments that involved monitoring incorporation of the thymidine analogue 5-ethynyl-2’deoxyuridine (EdU) into replicating DNA, and use of the tracker dye CellTrace Violet (CTV) to mark non-dividing parasites. Whether this represents long-term metabolic quiescence analogous to that in *T. gondii* and *Plasmodium spp.,* a slow-replicating phenotype as in *Leishmania spp.*, or temporary arrest induced by stress, as exhibited by *P. falciparum* and mammalian cells, is unresolved. In this latter context, the report that *T. cruzi* amastigotes have an intrinsic ability to reduce their replication rate by temporary cell cycle arrest in G1, as a response to stress, nutrient availability and drug treatment, may be of relevance (Dumoulin and Burleigh, 2018). It is not known whether these represent overlapping or distinct mechanisms for entering a quiescent state. This could have implications for drug design, immunological interventions, and our understanding of *T. cruzi* persistence.

Using highly sensitive bioluminescence and fluorescence imaging (Lewis *et al*. 2014; Costa *et al*. 2018; Ward *et al*. 2020), we demonstrated that the gastrointestinal tract, specifically the colon and stomach, is a key site of *T. cruzi* persistence during chronic murine infections. Smooth muscle myocytes in the circular muscle layer of the colonic gut wall are the predominant host cell type. In the chronic stage, the entire colon typically contains only a few hundred parasites, often concentrated in a small number of cells that can contain >100 parasites. During the acute stage, however, when the parasite burden is considerably higher and many cells are infected, host cells containing >50 parasites are rarely found. Persistent parasites are also frequently detected in the skin during chronic infections, and in C3H/CeN mice, the skeletal muscle (Lewis *et al*. 2016; Ward *et al*. 2020). Further studies have also shown that parasite replication is asynchronous in individual host cells, a process that is independent of tissue location or disease stage, that replication of the nuclear and mitochondrial genomes is non-coordinated within the intracellular population, and that replicating amastigotes and non-replicating trypomastigotes can co-exist in the same cell (Taylor *et al*. 2020).

We have developed tissue processing protocols and imaging procedures that allow us to routinely detect *T. cruzi* persistence foci during chronic murine infections at single cell resolution (Costa *et al*. 2018; Ward *et al*. 2020). Here, we describe experiments which provide new insights into parasite persistence, and demonstrate that chronic infections are associated with a reduced rate of parasite replication.

## Results

We sought to explore parasite replication by using CTV, a tracker dye that has been employed as a marker for spontaneous dormancy in *T. cruzi* amastigotes (Sánchez-Valdes *et al*. 2018). This succinimidyl ester dye diffuses into cells, binds covalently to the amine groups of proteins, and becomes fluorescent following cleavage by intracellular esterases (Filby *et al*. 2015). CTV fluorescence undergoes serial dilution with each round of parasite cell division, resulting in an inverse correlation between dye retention and proliferation rate. However, reports that CTV itself can inhibit cell division (Lacy Kamm *et al*. 2020) prompted us to first investigate toxicity towards *T. cruzi.* Trypomastigotes were labelled by incubation for 20 minutes in 5 or 10 µM CTV (Materials and methods), conditions that had been used previously to monitor parasite proliferation (Sánchez-Valdes *et al*. 2018). When these parasites were added to the Vero cell monolayer, we found they were 60% less infectious than trypomastigotes that had been incubated in the DMSO solvent alone (Figure 1A). In the first 48 hours post-infection, there was then limited division of intracellular CTV+ve parasites, with most trypanosomes in a state of growth arrest (Figure 1B and C). By 72 hours, replication had been more widely initiated, although the average number of amastigotes per infected cell was still significantly below that of the controls (Figure 1B). Microscopy also revealed heterogeneity in the intensity of CTV-staining within the *T. cruzi* population (Figure 1C and D), with many parasites failing to replicate, particularly in the first 36-48 hours post-infection. At lower CTV concentrations (1 and 2 µM), growth inhibition was less evident and fewer parasites retained the dye at 5 days post-infection (Figure 1D). Collectively, these experiments indicate that CTV is an inhibitor of trypomastigote infectivity and amastigote replication, and that the use of dye retention as a marker for dormancy and cell cycle arrest could lead to ambiguity. Furthermore, the heterogeneous nature of *T. cruzi* CTV-staining, even within individual host cells (Figure 1C and D, supplementary Video 1), could result in differential growth and development rates within the same intracellular parasite population.

**Figure 1.**
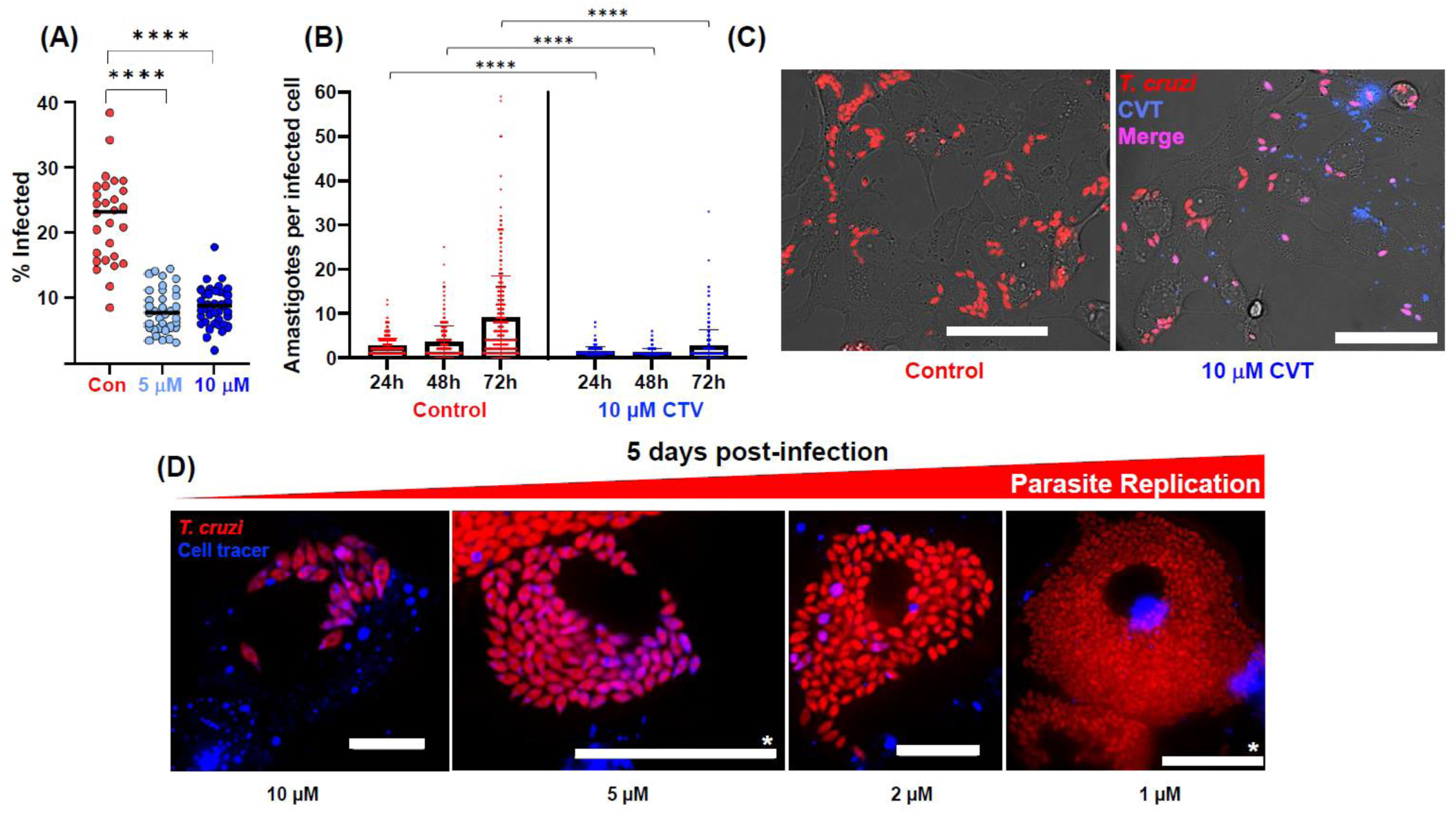
CellTrace Violet (CTV) reduces *T. cruzi* infectivity and inhibits intracellular proliferation. (A) *T. cruzi* CL-Luc::Scarlet trypomastigotes were incubated with either 5 or 10 µM CTV for 20 minutes and used to infect Vero cell monolayers at a multiplicity of infection of 10:1 (Materials and methods). 18 hours later, infection efficiency was determined by inspecting a total of 2,203 (control), 3,781 (5 µM), and 3,840 (10 µM) Vero cells. At least 300 of these were infected in each case. Each data point corresponds to a randomly acquired image and represents the mean percentage of cells infected. Differences between columns were analysed using a parametric one-way ANOVA with Tukey's post hoc pair wise comparisons. ****p=≤0.0001. (B) Vero cells infected with CTV−ve or CTV+ve trypomastigotes (as above) were incubated for the time periods indicated. The numbers of amastigotes per infected host cell were then determined by analysing >300 infected cells per treatment. Error bars represent the standard deviation from the mean. Data were analysed using a Wilcoxon rank sum test. (C) Images of Vero cells 36 hours after infection with CTV−ve or CTV+ve trypomastigotes. Red, fluorescent *T. cruzi* amastigotes. Fluorescent parasites containing the CTV tracer dye appear as purple on a red fluorescent background. Size bars=20 µM (D) Images of Vero cells 5 days after infection with trypomastigotes that had been incubated with various concentrations of CTV, as indicated. Blue, intracellular vesicles containing CVT. See also supplementary Video 1. Size bars=20 µM, except where indicated; *=50 µM.

Prior to infection, we removed non-bound CTV by quenching with addition of bovine serum (Materials and methods). Despite this, some dye was taken up by mammalian cells and retained in stained vesicles for several days (Figure 1C and D, supplementary Video 1). We found that it was important to ensure co-localisation of blue (CTV), red (fluorescent parasites) and/or green (DAPI - DNA) staining to avoid the risk of confusing amastigotes and spherical CTV-containing vesicles, both of which appear motile in the highly dynamic cytoplasmic environment.

We then used the thymidine analogue EdU to monitor parasite proliferation. In *T. cruzi*, incorporation of EdU provides a readout on the replicative status of both nuclear and mitochondrial DNA (kDNA) (Costa *et al*. 2018; Taylor *et al*. 2020). However, the procedure has to be used with caution, since EdU exposure *in vitro* can be associated with toxicity. In cultured mammalian cells, this is characterised by genome instability, DNA damage and cell cycle arrest (Kohlmeier *et al*. 2013; Zhao *et al*. 2013; Ligasová *et al*. 2015). Short-term exposure at lower concentrations (<12 hours, <10 µM) appears to have less impact, and does not perturb cell cycle kinetics (Pereira *et al*. 2017). Toxicity against *T. cruzi in vitro* has also been shown to be dependent on exposure time; 2 and 4 hours had only minor inhibitory effects on intracellular amastigotes, even at 70 µM, whereas with 24 hours continuous exposure, the IC^50^ dropped to 70 nM (Sykes *et al*. 2020). In contrast, when infected cells were cultured for 72 hours in presence of 100 µM EdU, there was no reported inhibition of amastigote replication (Sánchez-Valdéz *et al*. 2018). Given these conflicting observations, as a preliminary to *in vivo* studies, we assessed the *in vitro* kinetics and growth inhibitory effects of EdU on intracellular amastigotes of *T. cruzi* CL-Luc::Neon (a derivative of the CL Brener strain). This parasite reporter line expresses a fusion protein that is both bioluminescent and fluorescent (Costa *et al*. 2018). After only 10 minutes exposure, amastigotes were clearly labelled, and it was possible to distinguish those that were EdU+ve from those that were EdU-ve (Figure 2A). Similar heterogeneity, including differential labelling of nuclear and kDNA, was observed when infected cells were pulse labelled for 1 or 6 hours at different EdU concentrations (Figure 2B and C). This pattern results from asynchronous amastigote replication (Taylor *et al*. 2020), with EdU negativity/positivity determined by the position of individual parasites within the cell cycle during the period of exposure. When we assessed the extent of EdU growth inhibition after a 6-hour pulse and a 3-day chase period, we established an IC^50^ of 1.67µM, although the level of inhibition plateaued at 70% (Figure 2E). These outcomes are therefore consistent with those reported by Sykes *et al*. 2020.

**Figure 2.**
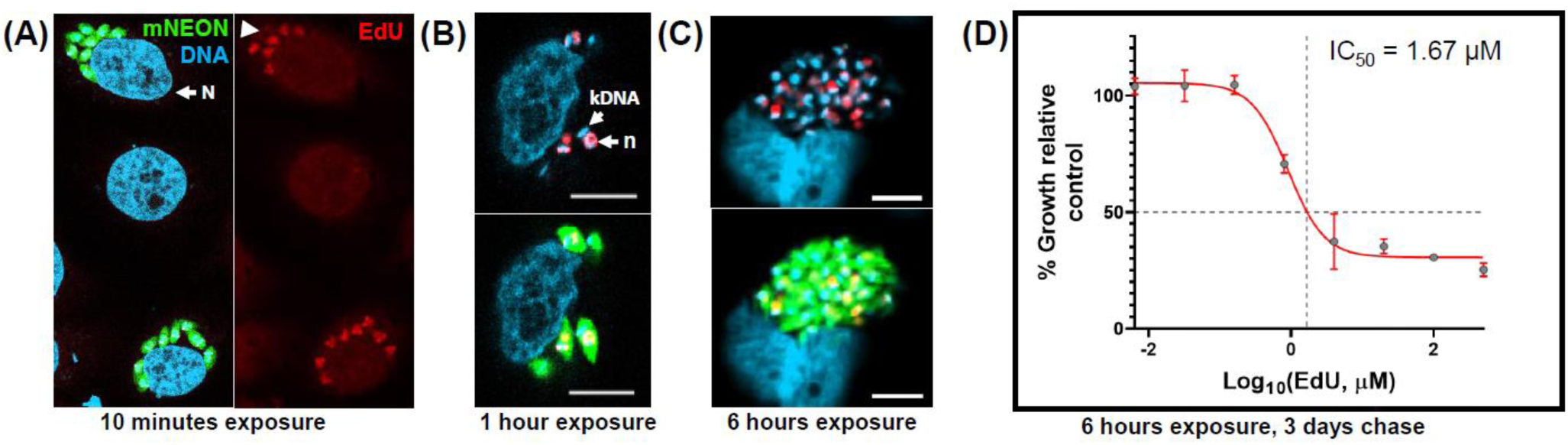
EdU incorporation by *T. cruzi* amastigotes *in vitro* is rapid, heterogeneous within the population, and can inhibit parasite growth. (A) *T. cruzi* CL-Luc::Neon trypomastigotes were used to infect MA104 cells at an MOI of 5:1 (Materials and methods). 2 days later, cultures were pulsed for 10 minutes with EdU (50 µM) and then examined by confocal microscopy. The image shows adjacent infected cells where in one instance, all 8 amastigotes are EdU+ve, whereas in the other, 2/8 are EdU−ve (indicated by white arrowhead). N, host cell nucleus. (B) EdU labelling of amastigotes after 1 hour exposure (40 µM). kDNA, kinetoplast DNA; n, parasite nucleus. (C) EdU labelling of amastigotes after 6 hours exposure (10 µM). Scale bars=10 µm in all cases. (D) MA104 cells were infected with trypomastigotes, and 2 days later, the cultures were pulsed for 6 hours with EdU at a range of concentrations, and then washed thoroughly. After a further 3 days incubation, amastigote growth was determined by assessing expression of the mNeonGreen reporter (Materials and methods). The inhibition curve was plotted using PRISM graphpad to establish the concentration of EdU that conferred 50% growth inhibition compared to untreated controls. Data were derived from 5 replicates (CI_95_ 0.98 µM-4.83 µM).

We next compared the dynamics of EdU incorporation by parasites during acute and chronic murine infections with the *T. cruzi* CL-Luc::Neon strain. The elimination half-life (T^1/2^) of EdU in mice has not been determined, but with other thymidine analogues, the period is relatively short. For example, the availability of bromodeoxyuridine (BrdU) for incorporation into mouse DNA is less than 15 minutes (Mandyam *et al*. 2007). Infected C3H/HeN mice were therefore given two EdU injections (each of 12.5 mg kg^-1^), 6 hours apart (Figure 3A), in an attempt to highlight a greater number of parasites where DNA replication was underway.

**Figure 3.**
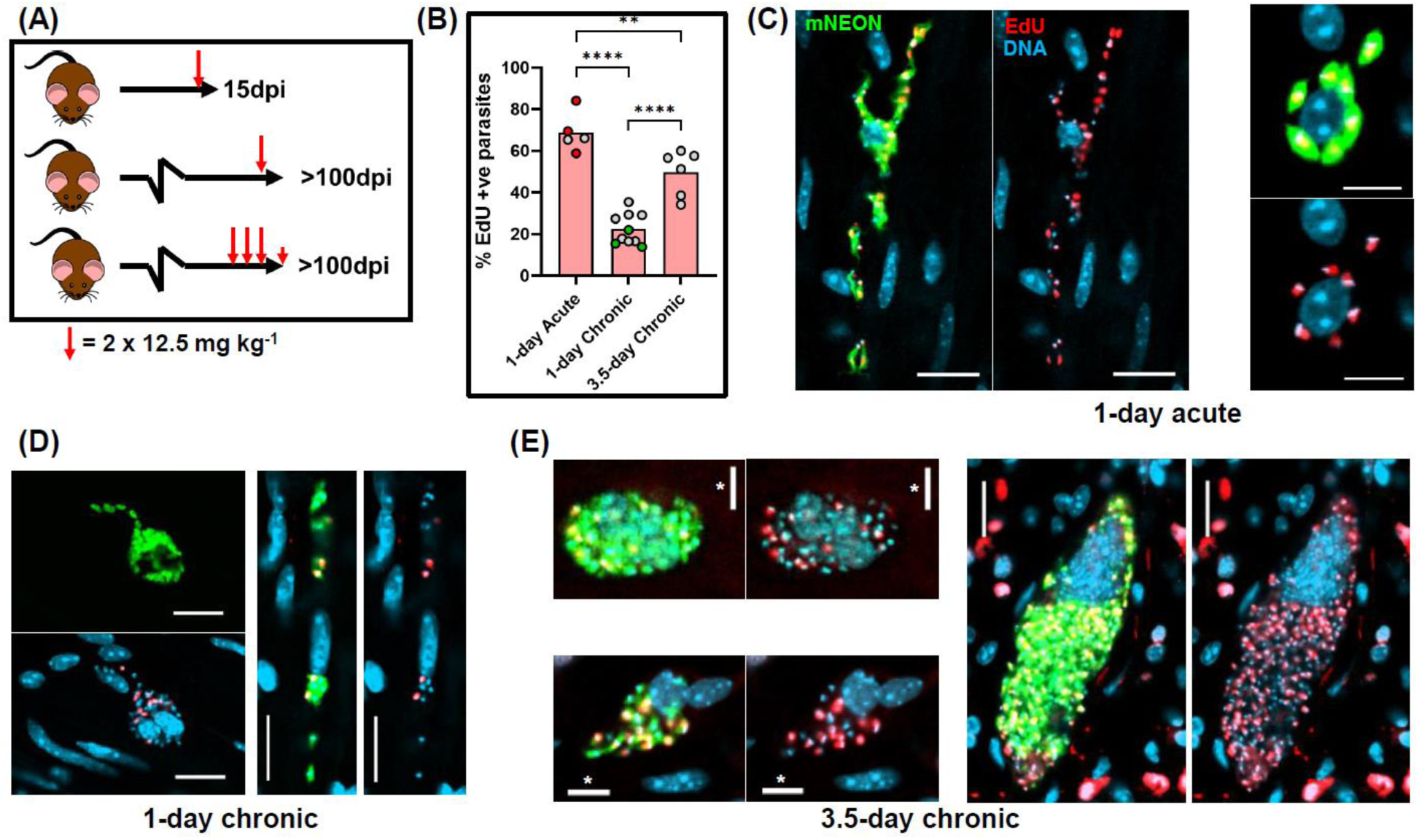
Intracellular *T. cruzi* replication, as inferred by EdU incorporation, is slower in the chronic stage than during acute infections. (A) Schematic showing experimental outline. C3H/HeN mice infected with *T. cruzi* CL-Luc::Neon were injected with EdU as indicated during the acute (15 days post-infection; dpi) or chronic stage (>100 dpi). Larger red arrows indicate 2 i.p. injections separated by 6 hours; smaller red arrow indicates a single i.p. injection on day 4. (B) Percentage of parasites that were EdU+ve under each treatment regimen. Colonic tissue was extracted from mice *post-mortem*, 18 hours after the second injection (1-day treatment) or 4 hours after the final injection (3.5-days treatment), and infected cells detected by *ex vivo* imaging and confocal microscopy (Materials and methods) (Figure 3 – figure supplement 1) (Ward *et al*. 2020). Each data point represents a single mouse. Red data points indicate mice aged ~150 days at the start of acute infection, and act as age matched controls for chronic stage mice. In the chronically infected mice, green data points indicate colons processed by standard histological sectioning, and grey points highlight those processed through peeling away of the mucosal layer and whole mounting the remaining colonic gut wall (Materials and methods). No significant differences were observed in the % EdU+ve parasites in colonic sections processed by each method (Wilcoxon rank sum test). For comparison of treatment conditions, statistical analysis was performed as described (Materials and methods); ****=p≤0.0001 and **=p≤0.01. (C) Representative images of infected colonic muscle cells from an acute stage mouse. Labelling: parasites, green; DNA, blue (DAPI staining); EdU, red. EdU labelling on a green background appears yellow. (D and E) Images of infected colonic muscle tissue from chronically infected mice after EdU labelling using the 1-day and 3.5-day regimen, respectively (see also Figure 3 – figure supplement 2). Scale bars=20 µm, except where indicated; *=10 µm.

At any one time, as judged by *in vitro* experiments, approximately 25-30% of amastigotes will be in S-phase (Dumoulin and Burleigh, 2018). In the majority of cases, the external gut wall mount methodology (Ward *et al.* 2020) was used to process the resulting tissue samples. The protocol enables the muscular coat, including the longitudinal and circular smooth muscle layers, which contain the majority of colon-localised parasites during chronic stage infections, to be visualised in their entirety at a 3-dimensional level with single-cell resolution (Materials and methods). Intracellular parasite numbers can be determined with accuracy by confocal microscopy using serial Z-stacking (Figure 3 – figure supplement 1). On occasions, infected host cells in colonic tissue were also investigated by coupling *ex vivo* bioluminescence-guided excision and confocal microscopy (Materials and methods) (Taylor *et al*. 2020).

Colon samples were excised *post-mortem*,18 hours after the second injection, and the incorporated EdU labelled by click chemistry. We observed significantly greater levels of parasite labelling in tissue obtained from acute stage mice than from those that were chronically infected (70% vs 20%) (Figure 3 and 4, Figure 3 – figure supplement 2). Therefore, during the acute stage, a greater fraction of the parasite population is replicating their nuclear and/or mitochondrial DNA at any specific point in time. By inference, the amastigote replication rate must be slower during chronic infections, at least in this tissue location. There were no differences in the data derived from colonic tissue processed by the two differing methodologies (grey and green dots, Figure 3B). We further observed that during the acute stage, there was a positive correlation between the number of parasites per infected cell and the percentage of parasites where DNA synthesis was ongoing (Figure 4B). Large parasites nests were less common during the acute stage, with few instances where infected cells contained more than 50 parasites (Figure 4A–C). We did attempt to quantify, for comparative purposes, the relative level of EdU labelling in each parasite. However, since most images were taken with whole mounted tissue sections, the variable depths of parasites from the surface made this technically challenging. As judged by visual inspection, the majority of EdU+ve parasites in any one cell were labelled to a similar extent (Figure 3C–E, Figure 3 – figure supplement 2). Since smooth muscle cells, the most frequently infected cell type in the colon (Ward *et al*. 2020), are typically non-dividing, the nuclei of host cells were generally unlabelled.

**Figure 4.**
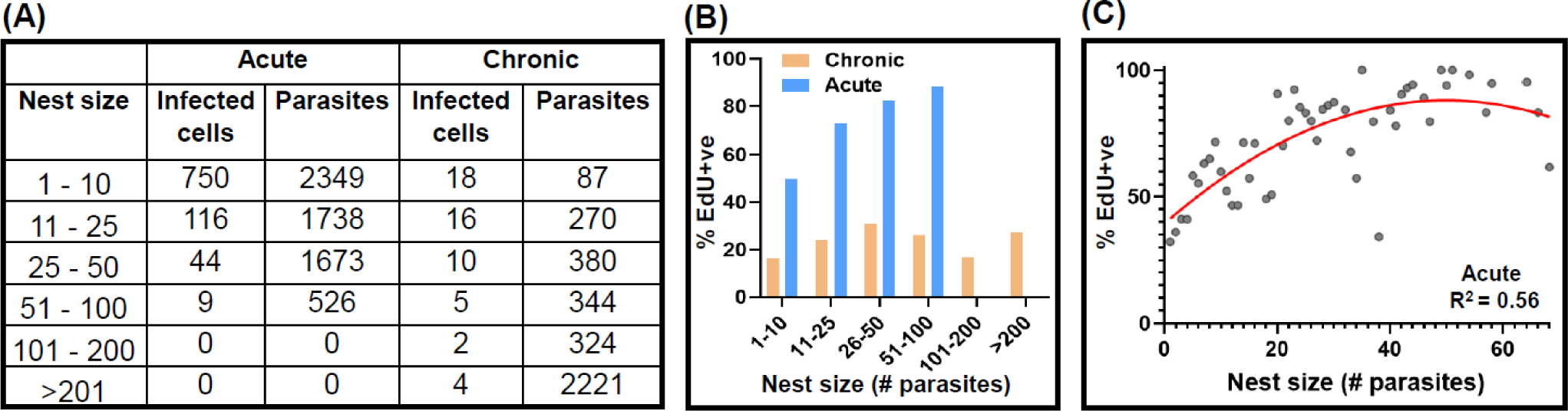
The inferred parasite replication rate is higher during the acute stage and correlates positively with nest size at this phase of the infection. (A) The number of infected cells (nests) detected in colonic gut wall tissue from mice in the acute (n=5) and chronic (n=7) stage, and the number of parasites found in each category. Tissue was processed using the colon peeling procedure (Materials and methods). (B) Percentage EdU+ve parasites in infected cells during acute and chronic infections in relation to nest size. (C) Relating nest size to the % EdU+ve parasites during acute stage infection. Each point corresponds to a specific nest size (x-axis), and the corresponding % EdU+ve mean percentage value across all mice (y-axis).

In chronically infected mice, only a minority of parasites incorporated EdU when the 1-day protocol was used (Figure 3 – figure supplement 2), and in ~20% of infected cells, none of the parasites were labelled (Figure 5B). A similar level of heterogeneous incorporation was observed in skeletal muscle, another site of parasite persistence in chronically infected C3H/HeN mice, and a tissue where parasites are often found in large nests (Figure 3 – figure supplement 3, as example). When labelling was extended over 3.5-days (a total of 7 injections) (Figure 3A), there was a 2.3-fold increase in the number of labelled parasites, with approximately half of those in the colon being EdU+ve (Figure 3B and 5C, Figure 3 – figure supplement 2). With this more prolonged protocol, every infected host cell that we examined contained at least one labelled parasite (Figure 5C). However, the percentage of EdU+ve parasites within the population was still significantly lower than during the acute stage, when the 1-day labelling protocol was used (Figure 3). In combination, these data indicate that during chronic infection of the colon, there is a general slowdown in the rate of parasite replication. As judged by bioluminescence *ex vivo* imaging of organs and tissues, neither the 1-day nor the 3.5-day EdU injection protocols had any detectable effect on the levels of infection or on tissue-specific parasite dissemination (Figure 3 – figure supplement 4).

**Figure 5.**
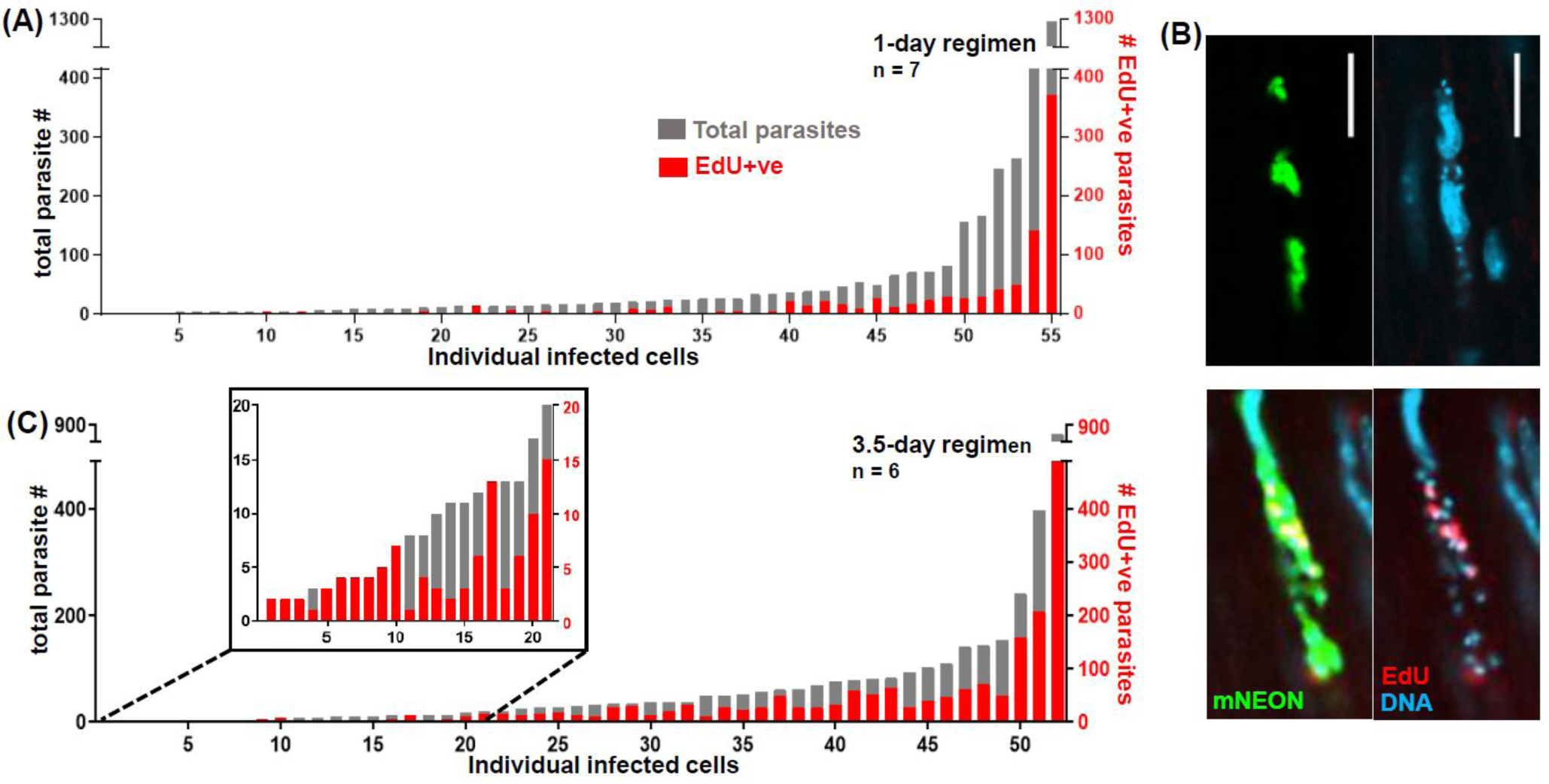
Increased chronic stage EdU incorporation by parasites with the 3.5-day regimen. (A) Level of EdU incorporation in the 55 infected cells detected in the whole mounted colonic gut walls from 7 chronic stage mice treated with the 1-day labelling regimen. The total parasite content (grey) and the number that were EdU+ve (red) are indicated. (B) Upper images; an infected cell containing no EdU+ve parasites after labelling with the 1-day protocol. Lower images; an infected cell from the same colonic tissue that contained EdU+ve parasites (see also figure supplement 3). Parasites, green. (C) EdU incorporation in 52 infected cells detected in 6 chronically infected mice treated with the 3.5-day labelling protocol. Every infected cell in the colonic gut walls of these mice contained at least 1 EdU+ve parasite.

To further investigate parasite replication during chronic stage infections, we undertook pulse chase experiments to assess the extent and stability of parasite labelling 7 and 14 days after EdU injection using the 1-day pulse protocol (Figure 6A). When mice were examined by *ex vivo* bioluminescence imaging after 14 days, there had been no measurable impact on the parasite burden or tissue distribution (Figure 3 – figure supplement 4). At the 7-day chase point, EdU+ve parasites were still readily detectable, although there was a 4-fold decrease in their relative abundance within the population, and only ~40% of infected cells contained any labelled parasites (Figure 6C, E and F). By 14 days post-injection, out of the 87 infected cells detected in the colons of 8 mice, just 1 contained EdU+ve amastigotes (Figure 6D, E and G). Using serial Z-stacking, we established that this infected cell contained 82 parasites, 42 of which were labelled. Given this profile, the most likely explanation is that this host cell had remained infected for at least 14 days, with the parasites in a state of low proliferation. The EdU+ve parasites cannot have undergone many replication cycles during this period, since the labelling intensity was similar to that in parasites where the chase period was only 1 day. Furthermore, in dividing cells, incorporated nucleosides become undetectable after 2 to 5 generations, assuming random segregation of daughter chromosomes (Kiel *et al*. 2007; Ganusov and De Boer RJ, 2013). It can be further inferred from the rarity of cells containing EdU+ve parasites (Figure 6D and G), that long-term occupancy of individual colonic smooth muscle cells by *T. cruzi* is not a common feature of chronic stage infections, even though this tissue is a site of parasite persistence. In the vast majority of cases therefore, the normal infection cycle of parasite replication, host cell lysis, and re-infection appears to continue during the chronic stage, albeit at a reduced rate. Finally, the observation of multiple labelled amastigotes within a single host cell 14 days after injection (Figure 6D) demonstrates that EdU is stable once it has been incorporated into the *T. cruzi* genome and that it is not susceptible to removal by metabolic or DNA repair pathways. This stability has similarities with the situation in mice, where Merkel cells labelled during pregnancy remained EdU+ve in off-spring 9 months after birth (Wright *et al*. 2017).

**Figure 6.**
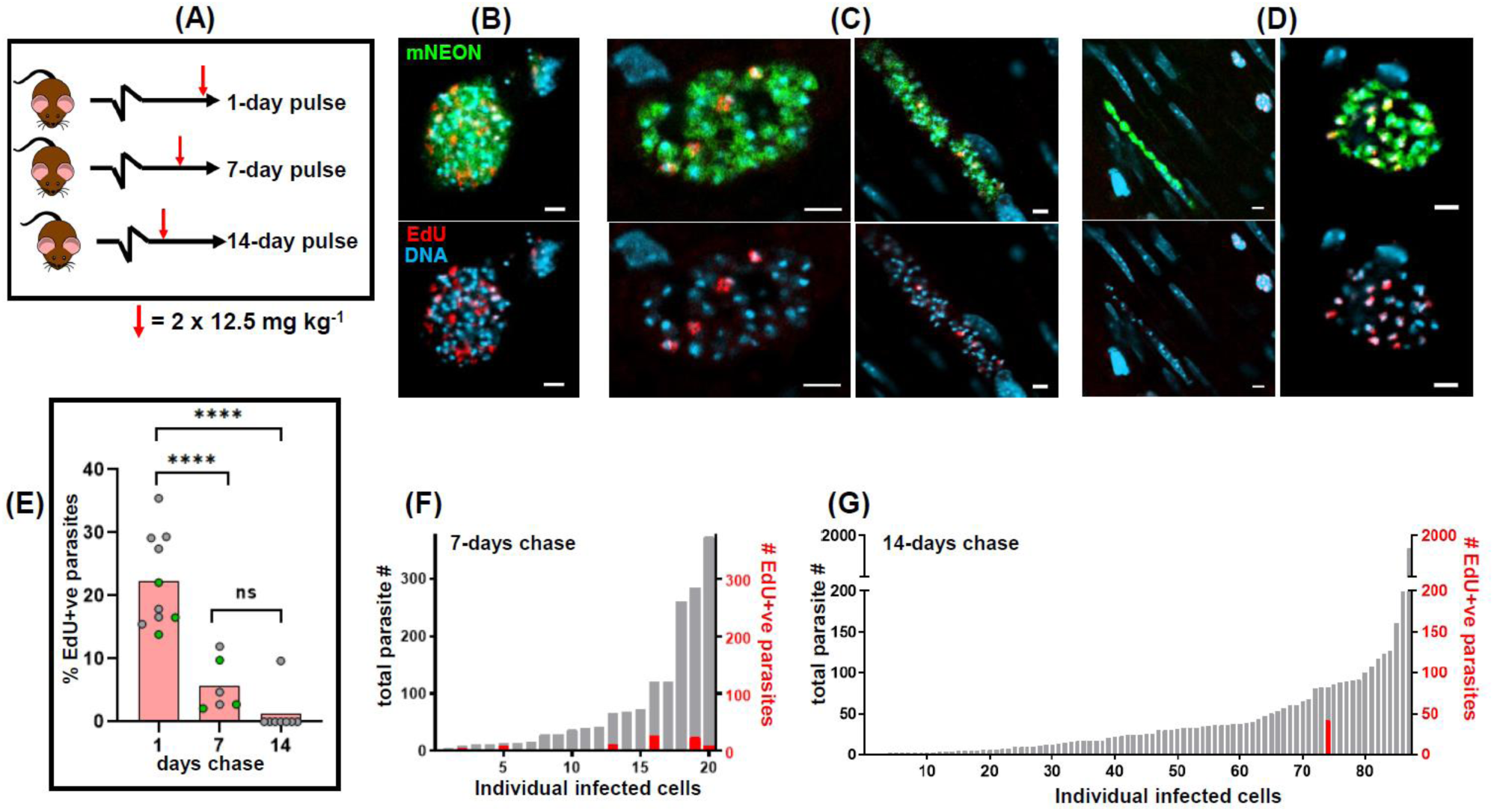
EdU pulse-chase experiments reveal that parasite occupation of individual colonic host cells during chronic stage infections is not long-term. (A) Schematic of the pulse-chase protocol. EdU was injected in two 12.5 mg kg^-1^ doses, 6 hours apart (as in Figure 3A). After 1, 7, or 14 days, mice were sacrificed, colonic tissue excised, and infected cells detected by *ex vivo* imaging and confocal microscopy (Materials and methods). (B) Representative images of parasite EdU incorporation following the 1-day chase regimen (see also Figure 3D). (C) EdU incorporation following a 7-day chase. (D) Left-hand image; EdU incorporation following a 14-day chase. Typically, infected cells contained no EdU+ve parasites. Right-hand image; the single example of an infected host containing EdU+ve parasites after an exhaustive search of colon mounts from 8 mice. Scale bars=20 µm. (E) Mean % EdU+ve parasites found in infected colonic cells. Each data point represents a single mouse; 1-day chase, n=10; 7-day chase, n=6; 14-day chase, n=8. The total number of parasites detected, imaged and designated as EdU+ve or EdU-ve in each mouse varied from 47 to 2468, with an average of 608. The green data points indicate colons processed by standard histological sectioning, and grey data points highlight those processed through peeling away of the mucosal layer and whole mounting of the remaining colonic gut wall (Materials and methods). Statistical analysis of treatment conditions was performed as described (Materials and methods); ****=p≤0.0001. There was no significant difference between the 7- and 14-day chase groups. (F) Percentage parasites that were EdU+ve in infected colonic cells from mice sacrificed 7 days post EdU pulse (n=3). Grey bar indicates total parasite number in each infected cell; red bar indicates the percentage EdU+ve. (G) Similar analysis of EdU positivity in infected cells found in mice 14 days post EdU pulse (n=8).

## Discussion

The report that *T. cruzi* can undergo a form of spontaneous dormancy has highlighted the possibility that the proliferation status of the parasite could have a role in long-term persistence and contribute to the high rate of treatment failure (Sánchez-Valdéz *et al*. 2018). Improvements in tissue processing and imaging procedures (Ward *et al*. 2020) have allowed us to explore this further by providing a platform to investigate parasite replication in the colon of chronically infected mice, a tissue that supports long-term *T. cruzi* persistence at extremely low levels (Lewis *et al*. 2014). Our major finding is that during chronic infections, the proportion of intracellular parasites in S-phase is significantly lower than it is during the acute stage. In acute infections, 70% of parasites were EdU+ve after a 1-day pulse-chase, compared with 20% during the chronic stage (Figure 3 and 4, Figure 3 – figure supplement 2). This is unlikely to reflect reduced EdU uptake or bioavailability during chronic infections, as the staining intensity of individual EdU+ve parasites was similar during both stages of the disease, with the only apparent difference being the proportion of amastigotes that were positive. The most parsimonious explanation is that *T. cruzi* amastigotes proliferate at a slower rate in the chronic stage, at least in this tissue location, and were therefore less likely to be replicating their DNA during the period of EdU exposure. It is implicit from this that *T. cruzi* amastigotes have the capability of responding to environmental signals that are specific to chronic and/or acute stage disease, such as nutrient availability or indicators of the immune response. This is line with previous reports that amastigote replication and cell cycle kinetics can be subject to reversible stress-induced inhibition *in vitro* (Dumoulin and Burleigh, 2018). A response mechanism of this type could also account for the correlation between amastigote growth rate and nest size during acute infections (Figure 4).

The heterogeneous nature of labelling during the chronic stage, with many parasites being EdU-ve (Figure 3, Figure 3 – figure supplement 2), should not be interpreted as being indicative of spontaneous dormancy. Rather it provides further evidence that parasite replication within individual host cells is asynchronous (Taylor *et al*. 2020). The observation that some intracellular parasites do not incorporate EdU reflects that amastigotes exist in a range of replicative states within individual infected cells, an inference supported by the cumulative nature of EdU labelling (Figure 3, Figure 3 – figure supplement 2). There are several possible fates for parasites labelled with EdU during chronic stage infections, as outlined in Figure 7.

**Figure 7.**
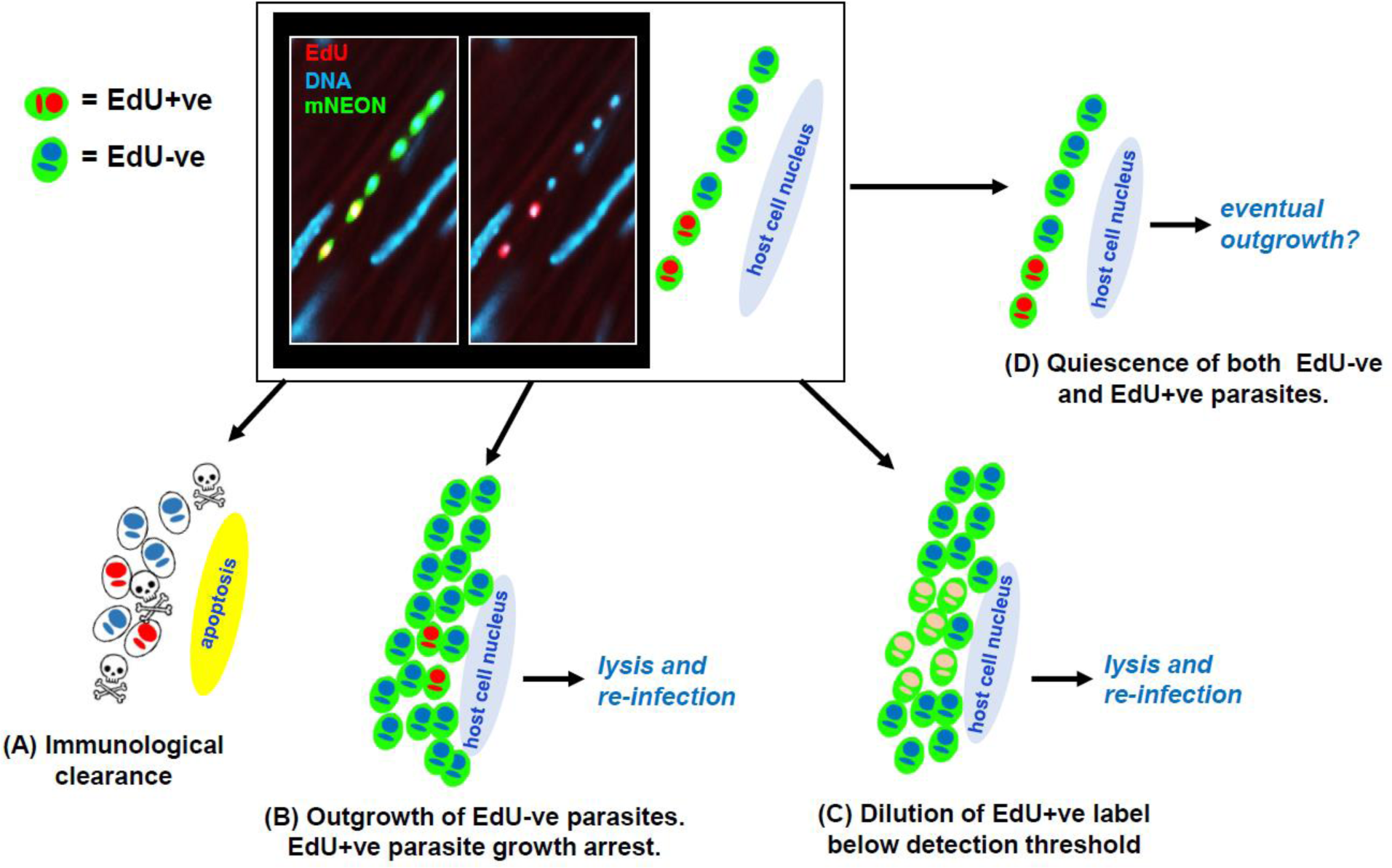
Schematic highlighting the possible fates of host cells and parasites following EdU labelling. The micrograph shows a colon smooth muscle section from a chronically infected mouse following treatment with the 1-day labelling regimen (Figure 3). EdU labelling (red) appears as yellow against the green background of parasite fluorescence. (A) Following the EdU pulse, the infected cell and parasites could be cleared by the host immune response. (B) There could be outgrowth of EdU-ve parasites in an infected cell. The EdU+ve subset, which would have been in S-phase during exposure, might enter cell cycle arrest after incorporation. (C) If EdU incorporation is below the toxicity threshold for cell cycle arrest, amastigote proliferation will lead to serial dilution of the label. (D) After EdU incorporation by a subset of parasites, all the amastigotes in the cell might enter a slow proliferative state. This appears to be a rare event (see Figure 6D and G).

Although incorporation of EdU into replicating DNA is widely used in proliferation studies, it can lead to a DNA damage response and cell cycle arrest, (Kohlmeier *et al*. 2013; Zhao *et al*. 2013; Ligasová *et al*. 2015). With *T. cruzi,* the measurable effect of EdU toxicity is time-dependent (Sykes *et al*. 2020), and here we showed that 6 hours exposure *in vitro* at 1-2 µM is sufficient to inhibit amastigote replication (Figure 2D). Therefore, studies on *T. cruzi* proliferation, dormancy and persistence could be confounded if EdU exposure is continuous. In the case of *in vivo* administration, EdU toxicity should be less problematic, because of the short clearance time of thymidine analogues (Mandyam *et al*. 2007). Consistent with this, we found no detectable impact of EdU exposure on parasite burden or tissue tropism in chronically infected mice (Figure 3 – figure supplement 4). In these infection experiments, where incorporation was employed as an end-point assay, providing snapshots of DNA replication within the parasite population, EdU toxicity would not be expected to compromise the outcome. This contrasts with *in vitro* experiments where EdU exposure can be continuous, resulting in interstrand cross-linking and double-strand breaks, which trigger DNA damage signalling and cell cycle arrest (Zhao *et al*. 2013). This is a ubiquitous response in all cells with DNA damage sensing machinery (Bielak-Zmijewska *et al*. 2018, Prokhorova *et al*. 2020). Spontaneous dormancy has been proposed as a mechanism that could account for parasite persistence after therapy (Sánchez-Valdes *et al*. 2018; Resende *et al*. 2020). However, in *T. cruzi,* the front-line drug benznidazole can cause mutagenesis, disruption to DNA-repair pathways, and chromosome instability (Rajão *et al*. 2014; Campos *et al*. 2017). Therefore, an alternative explanation could be that benznidazole-induced DNA damage responses trigger cell cycle arrest and a transient dormant-like state, which protects some parasites from further drug-induced toxicity, ultimately leading to relapse after the successful completion of DNA repair.

Our findings do not exclude the possibility that some parasites might have the potential to enter a canonical dormant state at specific points in the life cycle. However, they more strongly suggest, that rather than being a discrete biological stage, a dormancy-like phenotype in *T. cruzi* might be better described as representing one end of the normal proliferation spectrum. The cell cycle plasticity necessary for this has already been reported in amastigotes (Dumoulin and Burleigh, 2018). Therefore, the reduced rate of *T. cruzi* replication during the chronic stage could be a phenomenon more analogous to the partial biochemical quiescence and reduced proliferation exhibited by *Leishmania* (Kloehn *et al*. 2015; Mandell and Beverley, 2017), than to the more definitive quiescent state displayed by *T. gondii* and some *Plasmodium* species (Barrett *et al*. 2019). Resolving this question and understanding the mechanisms involved has particular importance for Chagas disease drug development strategies. As described in this paper (Figures 1 and 2), there are limitations to the cell tracker dye and DNA labelling methodologies that have previously been applied to investigate *T. cruzi* proliferation and quiescence. Therefore, new approaches are urgently required.

## Materials and methods

### Ethics statement

Animal work was performed under UK Home Office project licenses (PPL 70/8207 and P9AEE04E4) and approved by the LSHTM Animal Welfare and Ethical Review Board. All procedures were conducted in accordance with the UK Animals (Scientific Procedures) Act 1986 (ASPA).

### Parasites, mice and cell lines

The *T. cruzi* bioluminescent:fluorescent lines CL-Luc::mNeon or CL-Luc::Scarlet (Costa *et al.* 2018) were used throughout. Epimastigotes were grown in RPMI-1640, supplemented with 10% foetal bovine serum (FBS, BioSera), hemin (17 µg ml^-1^), trypticase (4.2 mg ml^-1^), penicillin (100 U ml^-1^) and streptomycin (100 μg ml^-1^), at 28°C. Metacyclic trypomastigotes were generated by culturing epimastigotes to stationary phase. *In vitro* studies were performed with MA104 and Vero African green monkey kidney cell lines. *In vivo* experiments used female C3H/HeN mice, initially aged 8-12 weeks, purchased from Charles Jackson (UK). Mice were maintained under specific pathogen-free conditions in individually ventilated cages. They experienced a 12-hour light/dark cycle and had access to food and water *ad libitum*.

### CellTrace *in vitro* assay

*T. cruzi* trypomastigotes were isolated by centrifugation and allowed to recover for 2 hours at 37ºC in high-glucose DMEM medium with 10% FBS, and then labelled with the CellTrace Violet (CTV) fluorescent dye (Thermo Fisher Scientific) according to the manufacturer’s protocol. Briefly, 2×10^6^ trypomastigotes were washed in PBS and then incubated for 20 minutes at 37°C in 10, 5, 2, or 1 µM CTV, protected from light. Unbound dye was quenched by the addition of 1 volume FBS and incubation for 5 minutes at 37°C. After washing (x2) in fresh complete medium, trypomastigotes were used for infection. Vero cells maintained in RPMI 10% FBS were trypsinized and seeded at 10^4^ or 10^5^ cells per well in 24-well plates containing cover slips, or in 8 well Ibidi µ-slides with a polymer coverslip, and allowed to attach for 6 hours before infection. Trypomastigotes were added at a multiplicity of infection of 10:1 (parasite:cell) and allowed to invade overnight (16-18 hours). Cultures were washed with PBS (x3) to remove non-invading parasites, and infected cultures incubated in RPMI with 2% FCS. Coverslips were fixed at different timepoints by transfer into a plate containing 4% paraformaldeyde for 30 minutes, then stained and mounted using Vectashield^®^ with DAPI, or with propidium iodide following RNase treatment.

Images and videos were acquired using an inverted Nikon Eclipse microscope. The slide containing the infected cells was moved along the x-y plane through a 580 nm LED illumination. Images and videos were collected using a 16-bit, 1-megapixel Pike AVT (F-100B) CCD camera set in the detector plane. An Olympus LMPlanFLN 40x/1.20 objective was used to collect the exit wave leaving the specimen. Time-lapse imaging was performed by placing the chamber slide on the microscope surrounded by an environmental chamber (OKOLab cage incubator) maintaining the cells and the microscope at 37°C and 5% CO^2^. Video projections and Z-stack sequences were created using the deconvolution app in the Nikon imaging software.

### *In vitro* parasite culturing and EdU labelling

Tissue culture trypomastigotes (TCTs) were derived after infecting MA104 cells with metacyclic trypomastigotes. MA104 cells were cultured in Minimum Essential Medium Eagle (MEM, Sigma-Aldrich.), supplemented with 5% FBS at 37°C, in 5% CO_2_. 24-well plates containing cover slips were seeded with 10^5^ cells per well and left for 48 hours. After reaching 95-100% confluency, they were infected with TCTs at a multiplicity of infection (MOI) of 5:1 (parasite:host cell). 18 hours later, external parasites were removed by washing (x3), fresh supplemented MEM was added, and the infections allowed to proceed.

EdU (Sigma-Aldrich) in PBS was diluted to the appropriate concentration in supplemented MEM (legend to Figure 2). The medium was removed, and the infected monolayer washed (x2), and fresh medium including EdU added. After the appropriate incubation period, cells were washed (x3). For EdU toxicity studies, parasite growth in infected cells was assessed in 96-well plates, 3 days after EdU addition, by measuring mNeonGreen fluorescence in a FLUOstar Omega plate reader (BMG LABTECH). Background fluorescence was calculated using uninfected MA104 cells (n=6). For microscopy, cells in the 24-well plates were washed (x2) and incubated for 45 minutes in 4% paraformaldehyde diluted in PBS. Cover slips were then removed and washed (x2) in PBS. EdU incorporation was assessed using a Click-iT Plus EdU AlexaFluor 555 Imaging kit (Invitrogen), as per manufacturer’s instructions, followed by washing (x2) with PBS, with coverslips then mounted in Vectashield. To allow precise counting of amastigotes, cells were imaged in 3-dimensions with a Zeiss LSM880 confocal microscope, using the Image Browser overlay function to add scale bars. Images were exported as .TIF files to generate figures.

### *In vivo* infections

CB17 SCID mice were infected with 1×10^4^ *T. cruzi* CL-Luc::mNeon TCTs and monitored by bioluminescence imaging (Lewis *et al*. 2015). At the peak of infection (~18 days), when bloodstream trypomastigotes were visible by microscopy, the mouse was euthanised (Taylor *et al*. 2019) and infected blood obtained by exsanguination. Trypomastigotes were washed in Dulbecco's Modified Eagle Medium, diluted to 5×10^3^ ml^-1^, and CH3/HeN mice injected i.p. with 1×10^3^ trypomastigotes.

### *In vivo* EdU labelling

The standard 1-day EdU treatment regimen involved two i.p. injections (12.5 mg kg^-1^ EdU in PBS) delivered 6 hours apart. The second injection took place 18 hours prior to sacrifice. For the 3.5-day treatment, the daily injection protocol (above) was extended for 3 days, with a final single injection on day 4, followed 4 hours later with euthanisation and necropsy. For acute stage experiments, mice were 14-16 days post-infection when EdU was administered, and for the chronic stage, mice had been infected for >100 days. Organs and tissues were subjected to *ex vivo* imaging, bioluminescent foci from skeletal muscle and the colon were excised, and processed for histology (Taylor *et al*. 2019). Where indicated, whole colons were removed from the gastrointestinal tract, pinned luminal side up, and the mucosal layer removed. Whole mounting of the entire external colonic gut wall was performed as described previously (Ward *et al*. 2020). Parasites were identified by mNeonGreen fluorescence using confocal microscopy, and carefully removed, together with ~5mm^2^ of surrounding tissue. Prior to a 2nd mounting, tissue pieces were processed for EdU detection by incubation overnight at 4°C in PBS containing 2.5% FBS and 0.5% triton-X (Sigma-Aldrich) and then washed in PBS (x2) (Taylor *et al*. 2020).

### Statistics

All statistical analyses were performed in GraphPad PRISM v8.0 and STATA v16.0., and the data expressed as the mean ± standard deviation of mean (SD), unless otherwise stated. *In vitro* EdU toxicity was calculated as % growth relative to non-treated controls. The data were fitted with a sigmoidal function with variable slope and the absolute IC^50^ value calculated by solving the function for X when Y= 50%. All data were tested for normality and homogeneity of variance using Shapiro-Wilk’s and Levene’s tests, respectively. Statistical comparisons between samples to analyse *in situ* EdU incorporation were performed using one-way ANOVA with post-hoc Tukey’s test for multiple comparisons. Data sets were analysed by non-parametric tests when variances were not homogenous. The % EdU incorporation in colon tissue sections vs. whole mount were compared using the Wilcoxon signed-rank test. The Kruskal-Wallis test was performed on the pulse-chase data. Statistical significance was accepted where p ≤ 0.05 (*=p ≤ 0.05, **=p ≤ 0.01, ***=p ≤ 0.001, ****p= ≤ 0.0001).

## Acknowlegements

This work was supported by UK Medical Research Council (MRC) Grant MR/T015969/1 to JMK and MRC LID (DTP) Studentship MR/N013638/1 to AIW.

---

**Figure 3 – figure supplement 1.**
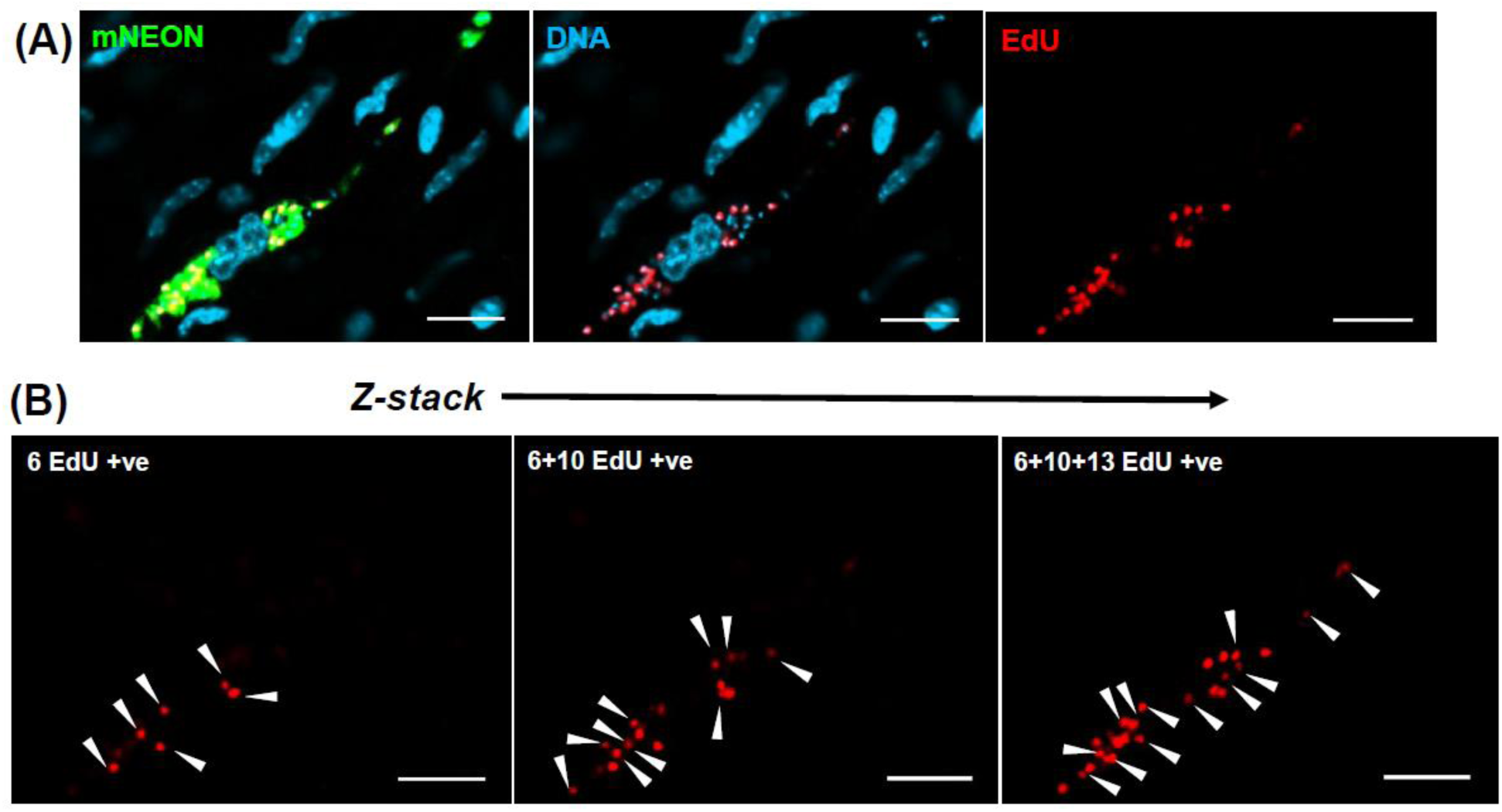
Determination of parasite numbers and EdU incorporation status in infected mouse cells using 3-dimensional confocal imaging. (A) *T. cruzi* infected cell in the colonic gut wall of a chronic stage C3H/HeN mouse, following treatment using the standard 3.5-day EdU labelling protocol (Figure 3A). All scale bars=20μm. (B) Assessment of EdU+ve parasites using a series of Z-stacked image slices from across the infected cell. 29 EdU+ve parasites were detected. The total number of parasites was determined in the same manner by visualising DAPI staining.

**Figure 3 – figure supplement 2.**
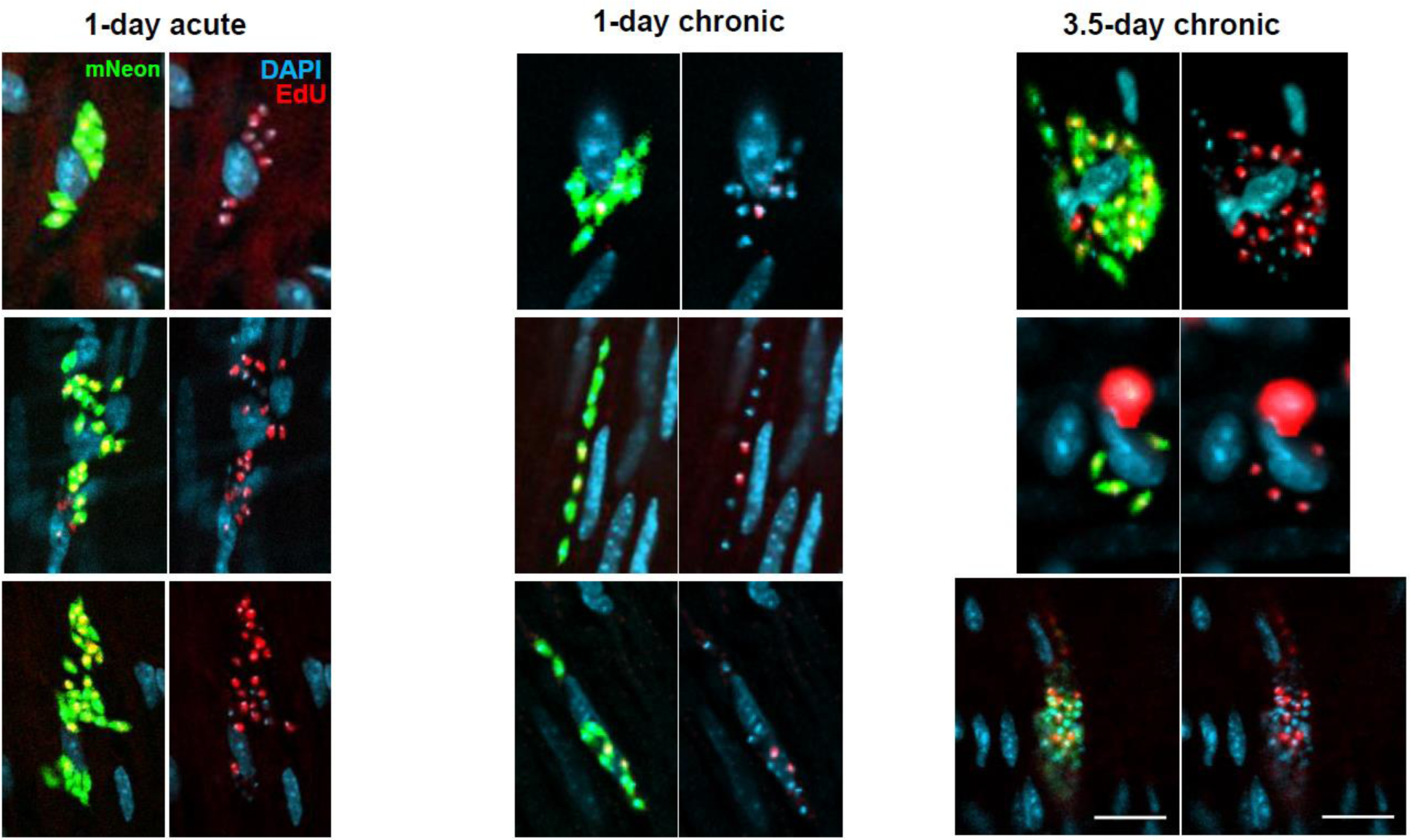
Further images demonstrating that, inferred from EdU incorporation, *T. cruzi* replication is slower in the chronic stage than in the acute (see also Figure 3). C3H/HeN mice infected with *T. cruzi* CL-Luc::Neon were injected with EdU during the acute or chronic stage. Colonic tissue was extracted from mice 18 hours after the second injection with 12.5 mg kg^-1^ EdU (1-day treatment, acute and chronic stage) or 4 hours after the final injection (3.5-days treatment, chronic stage), and infected cells detected by *ex vivo* imaging and confocal microscopy (Materials and methods). Labelling: parasites, green; DNA, blue (DAPI staining); EdU, red. EdU labelling on a green background appears yellow. For reference, scale bars=20 µm. These images form part of the data set collated to produce Figure 3B.

**Figure 3 – figure supplement 3.**
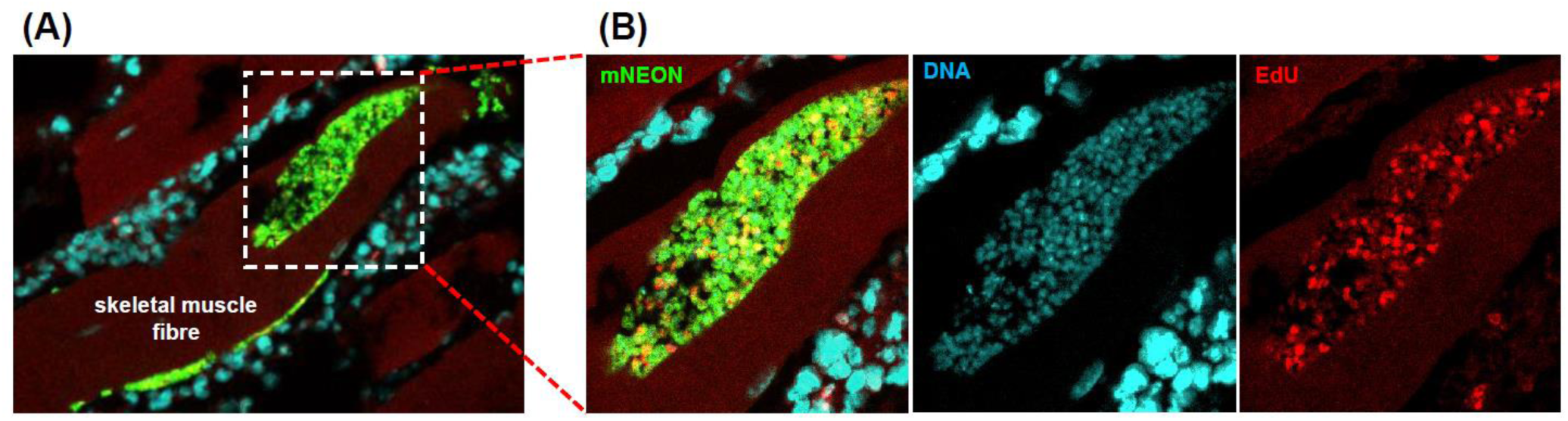
EdU incorporation by parasites in skeletal muscle during a chronic stage infection. (A) C3H/HeN mice, chronically infected with *T. cruzi* CL-Luc::Neon, were injected twice with EdU (12.5 mg kg^-1^; 6 hours apart). Skeletal muscle was excised 18 hours later and infected cells detected by *ex vivo* imaging and confocal microscopy (Materials and methods). (B) Enlarged images of nest showing fluorescent parasites (green), DNA staining (blue), and EdU incorporation (red).

**Figure 3 – supplement 4.**
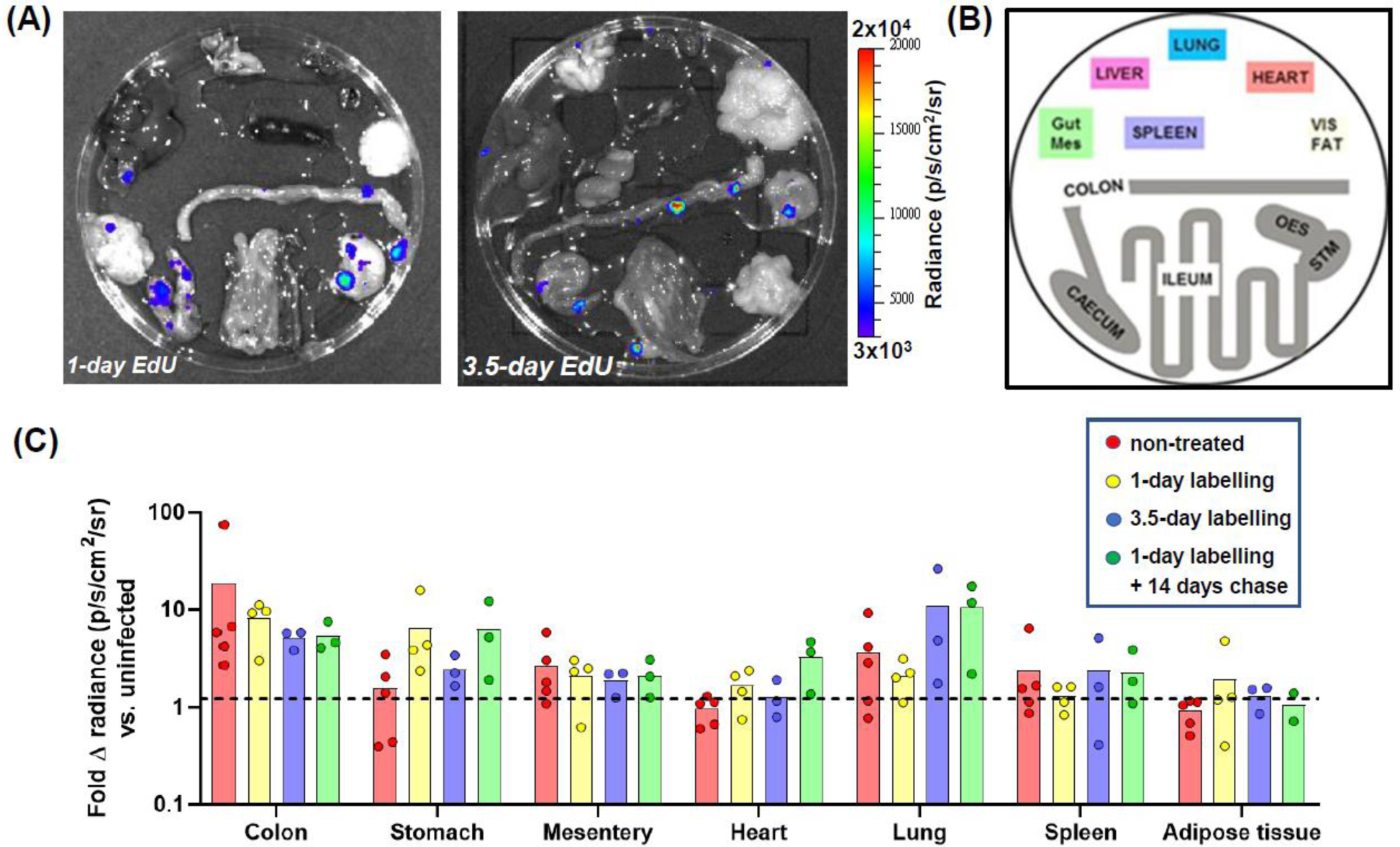
(A) *Ex vivo* bioluminescence imaging of organs and tissues from a C3H/HeN mouse chronically infected with *T. cruzi* CL-Luc::Neon, sacrificed after EdU labelling using the 1-day and 3.5-day protocol (Figure 3A). (B) Schematic showing the arrangement of organs. (C) Comparison of the fold increase in radiance (p/s/cm^2^/sr) above background associated with organs and tissues from chronically infected mice subjected to 1-day EdU labelling (n=4), 3.5-day EdU labelling (n=3), and 1-day EdU labelling, followed by a 14-day chase (n=3) (as in Figures 3 and 6). The dashed line indicates 2xSD above the background established from uninfected mice (n=4).

**Supplementary Video 1.**
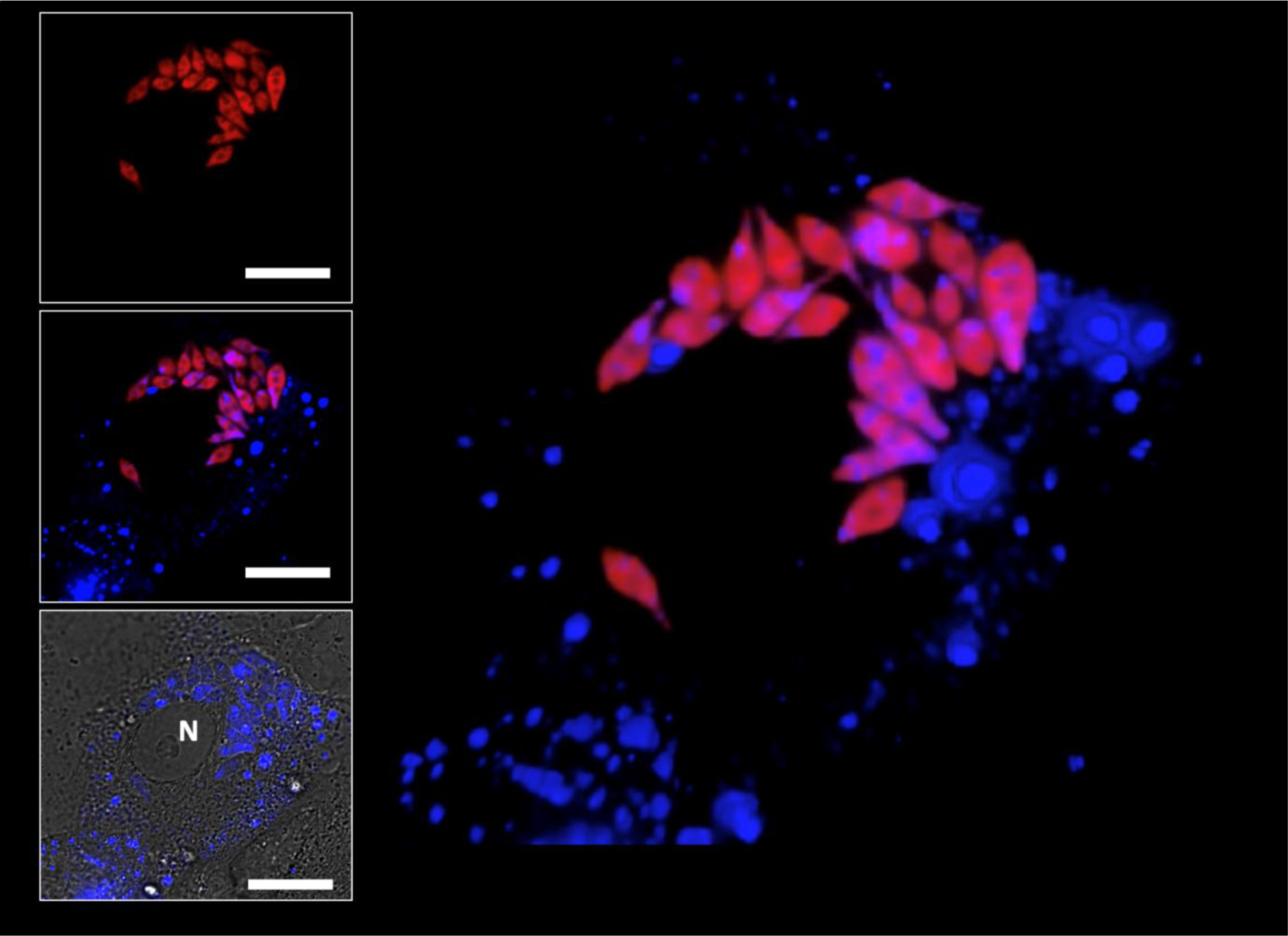
3-D video projection and rotation after Z-stack capture of a Vero cell 5 days post-infection with *T. cruzi* CLBr-Luc::Scarlet (red) (see also Figure 1D). Infective trypomastigotes had been pre-incubated with 10 µM CTV (Materials and methods). The asynchronous parasite population contains amastigotes displaying different sizes and shapes suggesting variability in their cell and/or developmental cycles. Within individual parasites, there was considerable heterogeneity in the intensity and location of CTV-staining. Similarly, in the host cell, CTV is widely dispersed (blue), frequently sequestered in vesicle-like structures that can be mistaken for amastigotes unless co-localising red fluorescence is confirmed. N=host cell nucleus. Size bars=20 µM. Video downloaded as separate file.

